# Modelling the role of immunity in reversion of viral antigenic sites

**DOI:** 10.1101/027995

**Authors:** Carmen H. S. Chan, Lloyd P. Sanders, Mark M. Tanaka

**Author notes:** Corresponding author *Email address:* (Carmen H. S. Chan).

## Abstract

Antigenic sites in viral pathogens exhibit distinctive evolutionary dynamics due to their role in evading recognition by host immunity. Antigenic selection is known to drive higher rates of non-synonymous substitution; less well understood is why differences are observed between viruses in their propensity to mutate to a novel or previously encountered amino acid. Here, we present a model to explain patterns of antigenic reversion and forward substitution in terms of the epidemiological and molecular processes of the viral population. We develop an analytical three-strain model and extend the analysis to a multi-site model to predict characteristics of observed sequence samples. Our model provides insight into how the balance between selection to escape immunity and to maintain viability is affected by the rate of mutational input. We also show that while low probabilities of reversion may be due to either a low cost of immune escape or slowly decaying host immunity, these two scenarios can be differentiated by the frequency patterns at antigenic sites. Comparison between frequency patterns of human influenza A (H3N2) and human RSV-A suggests that the increased rates of antigenic reversion in RSV-A is due to faster decaying immunity and not higher costs of escape.

## 1. Introduction

Viral evolution is shaped by both epidemiological effects on population dynamics, and molecular effects of mutations in the viral genome [1]. The combination of these effects generates distinctive dynamics at antigenic sites of viral proteins, which are the targets of host immune recognition. Selection for strains carrying antigenic changes that evade immune recognition result in elevated rates of non-synonymous substitution. It is unclear, however, why different dynamics of forward or reverse substitution are observed. Antigenic reversion has been reported frequently in viruses such as HIV [2, 3, 4], respiratory syncytial virus (RSV) [5] and hepatitis C [6, 7], and less frequently in other viruses such as influenza [8, 9], parvovirus [10], hepatitis A [11] and polio [12]. Various explanations for occurrence of reversion have been proposed, such as changing immunity [5], a limited antigenic repertoire [5, 9], or constraints on function [11, 7, 8], but it is not understood how the relative influence of these effects can generate differences in observed rates of reversion.

The difficulty in evaluating the contribution of selective mechanisms is due to the lack of methods that model both epidemiological and molecular dynamics. Phylodynamic approaches [13] incorporating epidemiological models into a coalescent framework have provided insight into the origins and spread of novel pathogens. However, they assume that molecular changes do not affect epidemiological dynamics, and are uninformative about selection. In contrast, codon-based approaches [14, 15] aim to identify sites that contribute to the adaptation of a virus, but they assume that the population size is constant and that the selection coefficient is constant at each site. Various modifications of the substitution model allow for different selective effects based on directionality or target residue [16, 2], but retain the assumption that substitution occurs as a time-homogeneous process which is not affected by population dynamics. To understand how the probability of reversion at antigenic sites is affected by both selective constraint against molecular changes and selection to evade immune recognition, there is a need to incorporate the time-dependence imposed by epidemiological dynamics into the substitution process.

Models of pathogen dynamics have shown that reversion probabilities are affected by fitness costs [17, 18, 3, 19], at both the within-host and between-host level, and the availability of susceptible hosts [3], at the between-host level. However, these models were developed in the context of HIV escape mutations. HIV infects host chronically, with host susceptibility determined by human leukocyte antigen (HLA) type, which does not vary over time. Due to these infection dynamics the prevalence of each strain changes relatively slowly, and is expected to eventually stabilise [3]. In contrast, for acute infections such as human influenza and RSV where transmission occurs frequently and host immunity can last for much longer than the duration of the infection, the structure of host immunity can vary rapidly over time. Due to differences in the dynamics of selection, we expect antigenic selection to have qualitatively different effects on sequence changes at antigenic sites compared to constant selective pressure [1].

Here, we examine the probability of antigenic reversion in an epidemiological model, which describes the complex ecology of multiple viral strains with cross-immunity competing for susceptible hosts. This model allows us to quantify the relative advantage of an antigenically novel mutation, compared to a reversion which may be antigenically less advantageous, but improves transmission. Using both a simple three-strain model and simulations with multiple codon sites, we examine the effect of the duration of host immunity, selective costs, population size, and the basic reproductive ratio. We show that these effects lead to distinctive dynamics in the frequencies of derived amino acids, which is informative about the duration of host immunity and strength of selective constraint. Time-structured sequence data from influenza and RSV are compared to simulated sequences, and we discuss what these results imply about the relative effects of host immunity and functional constraint.

## 2. Methods

### 2.1. Simple analytical model for antigenic reversion

The simplest model containing reversion is a system where the population has mutated away from the ancestral state, and potentially can mutate either back to the ancestral state (reversion) or to a novel state (forward substitution). In an epidemiological context, we consider a viral population described by a three-strain SIRS model [20]. The viral population is initially of strain 0 (ancestral state), which is then replaced with strain 1, and can subsequently be replaced by either strain 0 (reversion) or strain 2 (forward substitution).

We assume a large host population of constant size *N*, with homogeneous mixing, so that the dynamics of the number of hosts which are susceptible *S_i_,* infected *I_i_,* and recovered with immunity *R_i_,* for strains *i* = 0,1, 2 can be described by

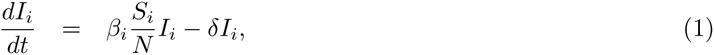

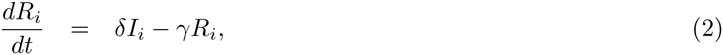

with transmission rate *β_i_*, recovery rate *δ*, and immunity that decays at rate *γ*. Interactions between strains are described by the implicitly defined term *S_i_*, which is the number of hosts susceptible to strain *i*. Assuming that each host can only be infected by a single strain at a time, and prior infection with strain *j* reduces susceptibility to strain *i* by a factor *σ_ij_*, the relationship between susceptible and immune hosts is given by

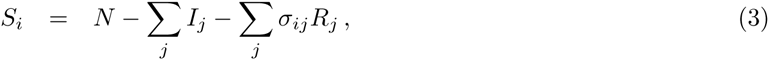

with the constraint that *S_i_* > 0 for any strain *i*. All uninfected hosts 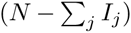 can be categorised as either susceptible (*S_i_*) or immune 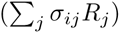 to strain *i*. The similarity between this model and the status-based model with polarised immunity developed by Gog and Grenfell [21] becomes evident when we differentiate Equation (3) to give

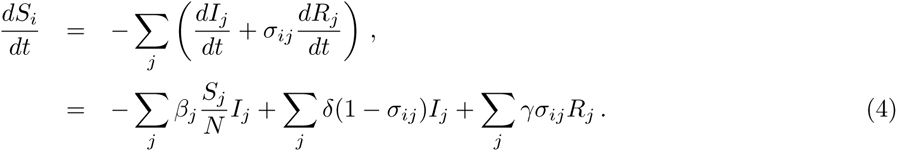

The main difference is that we retain the history of infections accumulated across the population through the additional set of variables, *R_i_*. This allows us to obtain analytical expressions for the number of hosts susceptible to all strains as functions of the same set of variables, as shown in Equation (3). In contrast to the Gog and Grenfell [21] model assuming polarised immunity, we assume a model of partial additive immunity. A host that was infected twice with strain *i* at times *t*_1_ and *t*_2_ will contribute 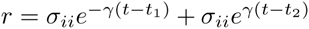 to *R_i_* at time *t*. This additive structure can be easily generalised to incorporate multiple strains. However, our model allows a host to contribute *r* > 1 after multiple re-infections, so we tend to inflate *R_i_*. The effect of this approximation is examined in greater detail for a single strain system in Appendix A.1. Overall, the effect of the approximation is to reduce *I*, but leave *S* unchanged. The approximation also tends to have minimal effect when *σ* is small, or when the rate of immune decay varies between hosts.

Our model of partial additive immunity generates similar dynamics to the Gog and Grenfell model [21]. From Equation (4), it can seen that hosts infected with strain *j* are removed from the susceptible class *S_i_*, and then a proportion 1 – *σ_ij_* of all infected hosts are returned to the susceptible class on recovery, so that the overall contribution of immunity is *σ_ij_β_j_S_j_I_j_/N*, which is similar to the *σ_ij_β_j_S_i_I_j_/N* term in the Gog and Grenfell [21] model. The difference in the *S_i_* and *S_j_* term arises because in the Gog and Grenfell [21] model, immunity arises from exposure, but in our model, immunity is only generated when infection occurs.

The strict exclusion of co-infection involves a second approximation, where an infection by any strain *j* will always be removed from *S_i_* but not from *R_i_* [first term in Equation (4)]. This occurs because while it is possible to distinguish between *S_i_* and *R_i_* at the time of infection from strain *j*, it is not possible, at the time of recovery from strain *j*, to determine whether the host was previously susceptible or immune to strain *i*. Our approximation leads to an underestimation of *S_i_*. We expect this to have a small effect as the bias lasts only for the duration of the infection. In addition, strains which are closest to the current circulating strain *j* will not be heavily affected (*σ_ij_* ≈ 1); the most heavily affected strains are those distant from strain *j* which are likely to be no longer circulating.

Using this model, we examine the effect of cross-immunity *σ_ij_*, immunity duration *γ* and selective costs incurred by antigenic escape *s*. The rate of immune decay *γ* includes the loss of immunity by the death and migration of immune hosts as well as the loss of immunity in individual hosts. The selective cost is parametrized through a reduction in the strain-specific transmission rate so that *β*_0_ = *β*, *β*_1_ = *β*(1 − *s*) and *β*_2_ = *β*(1 − *s*)^2^. To understand the effect of these parameters, we first characterise the number of susceptible hosts to each strain at equilibrium, and use this to determine probabilities of fixation, assuming a single strain appears at a time.

We assume the population is initially infected with only strain 0, which is maintained at equilibrium until strain 1 emerges at time *t*_1_. Strain 1, then replaces strain 0 and equilibrates until time *t*_2_, when a third strain (either strain 0 or strain 2) emerges and can potentially replace strain 1. These equilibrium assumptions allow us to characterise host immunity accumulated due to infection by strain 0 at *t*_1_ (denoted 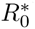), and host immunity accumulated due to infection by strain 1 at *t*_2_ (denoted 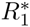), which then allows us to evaluate the probability of strain 0 or 2 emerging at time *t*_2_.

The equilibrium is obtained by setting the derivative of *S_i_* and *I_i_* to zero. When the viral population consists of only one strain, the endemic equilibrium, which is asymptotically, locally stable when the basic reproductive ratio *β_i_*/*δ* > 1 [20], is given by

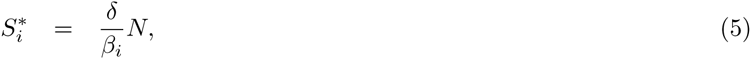

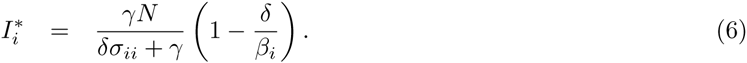

We assume that at time *t*_1_, when strain 1 emerges, the population remains close to equilibrium. As strain 1 has only just emerged and strain 2 has not yet occurred, the cross-immunity terms in Equation (3) can be ignored so that it contains only terms of subscript *i* = 0. Substitution of Equations (5) and (6) into Equation (3) gives

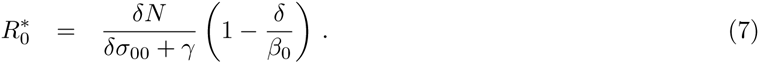

Now, consider a later time *t*_2_, when a third strain (either 0 or 2) emerges and can potentially replace strain 1. Again, we assume that strain 1 remains close to equilibrium and that the third strain has had negligible effect on immunity. In addition, we assume that immunity due to infection by strain 0 has decayed exponentially since time *t*_1_, so that Equation (3) can be approximated as

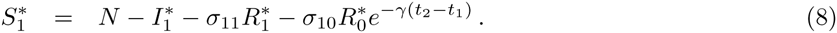

Substituting Equations (5) and (6) into (8) then gives

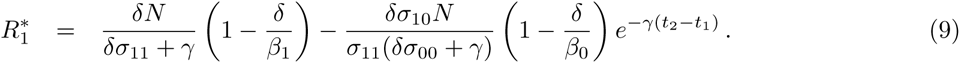

Having obtained an expression for 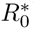 and 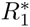, we can now compute the proportion of hosts that are susceptible to each strain, *p_i_*(*τ*) = *S_i_*(*τ*)/*N*, where *τ* = *t*_2_ − *t*_1_ is the time since the emergence of strain 1. Thus,

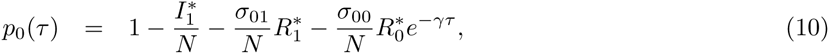

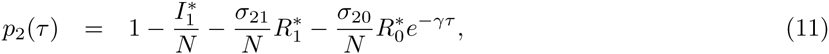

which can be written in the form

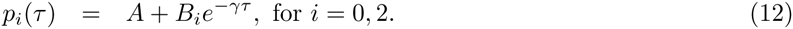

Assuming that cross-immunity is additive with respect to the number of antigenic differences (*σ_ii_* = *σ*, *σ*_01_ = *σ*_10_ = *σ*_21_ = *σ*/2 and *σ*_20_ = 0), the coefficients simplify to

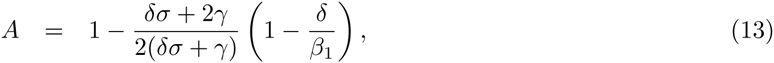

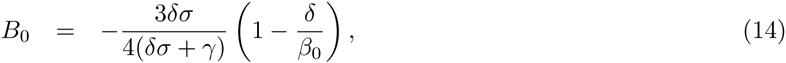

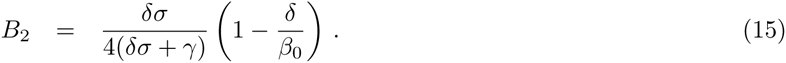

Note that we expect that prior immunity reduces infection against an unmutated strain at appreciable levels (*σ* ≫ 0.1) and that immunity lasts for much longer than the infection duration (*γ* ≪ *δ*). Within the parameter range of interest, the fractional terms containing *δ*, *σ* and *γ* in Equations (13-15) approach constants, so that *A* is approximately a function of only *β*_1_/*δ* and *B*_0_ and *B*_2_ are approximately functions of only *β*_0_/*δ*.

We calculate the probability of a strain generated by reversion or forward mutation at time *t*_2_ giving rise to a new epidemic by approximating the emergence of a new strain as a linear birth-death process. Ignoring initial changes in host susceptibility, the probability that a new strain reaches fixation [22] is given by

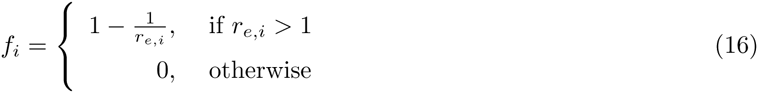

where *r_e,i_* = *β_i_p_i_*/*δ* denotes the effective reproductive ratio of the new strain *i* at the time of emergence. Using Equations (12-15), at time *τ* after strain 1 has reached equilibrium, we compute the probability of fixation for strain 0 (reversion) and strain 2 (forward substitution) to be

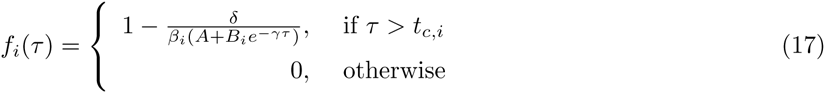

where the threshold *t_c,i_* is given by

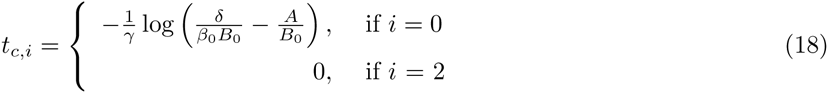

The probability of reversion given fixation is therefore

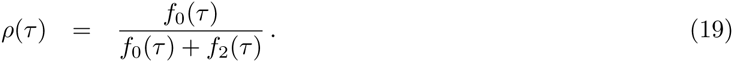

The probability of reversion is low immediately after the strain 0 has been replaced; in fact from Equations (17-19), it is zero for *τ* < *t_c_*,_0_. Asymptotically, if all prior immunity against strain 0 has decayed, then the exponential term in the denominator of Equation (17) approaches zero, thus giving

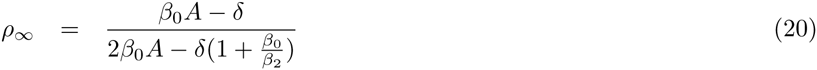

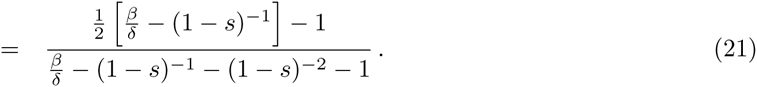

In summary, Equation (19) describes the combined effect of immunity *γ* and functional constraint *s* on the probability of reversion at some time *τ* after immunity has begun to wane from equilibrium levels. Whereas the long-term asymptote *ρ*_∞_, given by Equation (21), shows the effect of functional constraint in the absence of immunity.

### 2.2. Multi-site simulation model

To verify our theoretical model, and to examine the impact of increasing the antigenic space, we develop a stochastic computer simulation model where each infection is associated with a sequence of antigenic sites. Population dynamics are similar to the analytical model (see Table 1 for a complete list of parameters), but in the multi-site simulation, we explicitly model the mutation process. In the analytical model, we assumed the emergence of three strains at specified times, and calculated the probability that these strains would reach fixation. In contrast, for the simulation model, we allow mutations to occur stochastically at any antigenic site throughout the simulation; thus, new strains may emerge before the old strain reaches equilibrium and even favourable mutations may be lost due to stochasticity.

**Table 1:**
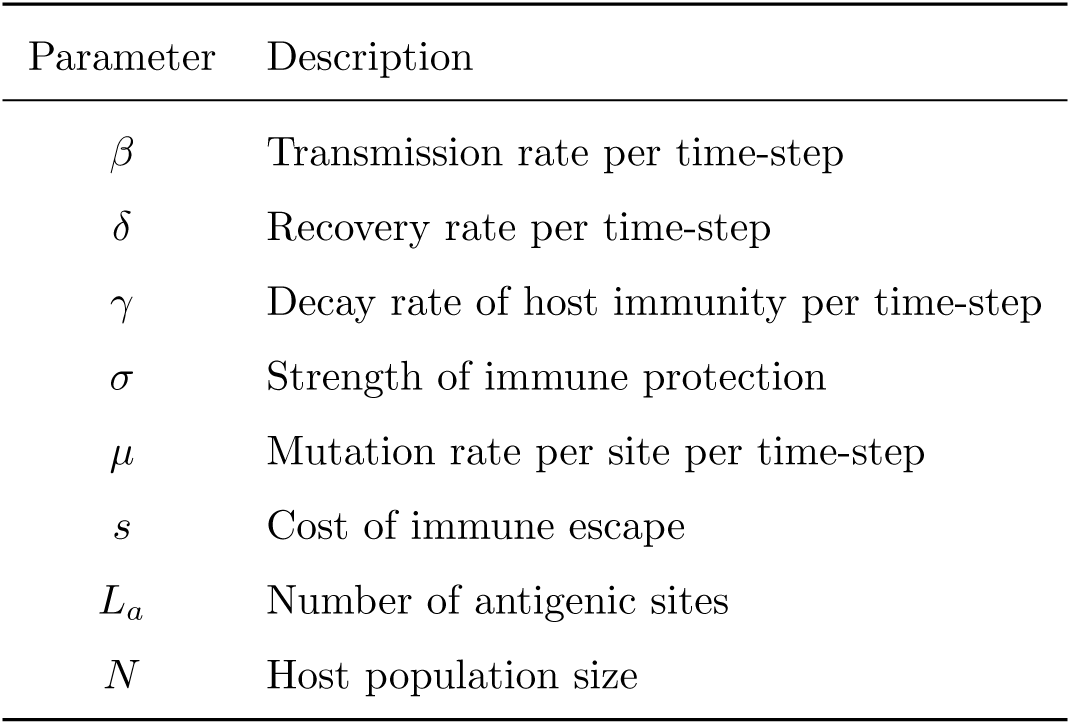
Table of parameters used in the multi-site simulation model.

We implement two models using different representations of the antigenic space. The first model uses a bit-string representation so that each of the *L_a_* antigenic sites can take values of **v** = {0,1}, and a change at any site away from the ancestral state (0) will reduce transmissibility. The bit-string model with two sites has a antigenic space similar to the analytical model. In the second model, we use a more realistic codon representation. Sites can mutate to any one of the 64 possible codons, but viral fitness is only affected by non-synonymous changes (i.e., **v** consists of the 20 amino acids). Specifically, any amino-acid change will affect cross-immunity, but only changes from the ancestral amino acid to a derived state will reduce transmissibility. The ancestral codon sequence is determined at the beginning of each simulation by randomly sampling *L_a_* non-terminating codons with uniform probability.

Throughout the simulation, we track the number of infected hosts *I*, the genotype of each infection, and the immune status of the host population. The last variable is stored in the immunity matrix consisting of 2 × *L_a_* elements for the bit-string model, or 20 × *L_a_* for the codon model, where each element *r_v,j_* stores the number of people with immunity to a value of *v* at site *j*. That is, *r_v,j_* stores the site-specific immunity accumulated across the whole population, and we compute the immunity against any viral genotype by summing across these values (described below).

The multi-site model is implemented as a discrete time simulation [22], with a time-step of one day. The system is initialised with a naive population ( *r_v,j_* = 0 for all *v* and *j*) and an infected host which carries the ancestral strain. At each time-step, the population changes according to SIRS dynamics, with the following events occurring:

1. Mutation: The number of mutations that occur in the viral population in each time-step is drawn from a Poisson distribution with mean *μIL_a_*, where *μ* is the mutation rate per site per time-step, and occur uniformly across all sites and all individuals. For the codon model, the probability of any codon occurring at the mutated site is specified by the Kimura two-parameter model [23] with a transition-transversion rate of *κ* = 3.
2. Transmission: The number of potential new infections which occur in each time-step is a Poisson random variable *X* ~ Pois(Λ), where 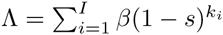 is the force of infection. The scaling factors 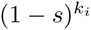 account for the reduction in transmission of genotype *i* due to the cost of *k_i_* changes away from the ancestral strain. The genotypes of the *X* potential infections are determined by multinomial sampling according to 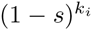, to account for variation in transmissibility within the viral population. We can then calculate the probability of each potential infection *i* encountering a susceptible host, given by

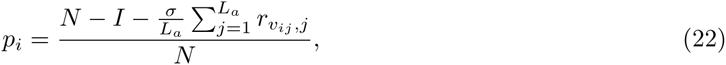

where *r_vij,j_* is the level of recognition against a particular antigenic site as described above. Equation (22) corresponds to Equations (10-11) in the analytical model. The success of the potential infection is determined using a Bernoulli random variable *U* ~ Bernoulli(*p_i_*). If *U* = 1, a new infection is generated with a genotype identical to the parent.
3. Recovery: The number of infected hosts which recover in each time-step is Poisson with mean *δI* truncated with an upper bound of *I* − 1. Each recovered host *i* is drawn from the infected population with uniform probability and increases immunity to allele *v_ij_* at site *j* = 1, …, *L_a_*. That is, for each recovery, we update *L_a_* elements of the immunity matrix

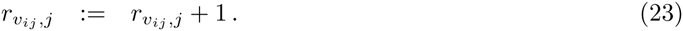
4. Decay of host immunity (across the whole population) is simulated by reducing *r_v,j_* for all antigenic states *v* ∈ **v** at each site *j* = 1, …, *L_a_* by a binomial random variable,

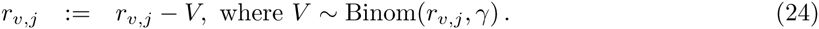

Note that the epidemic is artificially prevented from extinction. The forcing mechanism is necessary as we have, for simplicity, not included a migration term. In stochastic models of recurrent epidemics, the infection frequently dies out without re-introduction by migration, particularly in smaller populations [24, 25].

## 3. Results

Using the analytical and simulation models, we examine how the epidemiology of the virus affects the probability of reversion at antigenic sites. We first describe the dynamics of the simple three-strain model (Section 3.1), before examining the time dependence of this system (Section 3.2) and the effect of the epidemiological parameters (Section 3.3). The combined effect of these interacting factors on the observed amino acid frequencies is described in Section 3.4, and we compare this to sequence data for human influenza A (H3N2) and RSV-A in Section 3.5.

### 3.1. Dynamics of changing susceptibility

To provide some intuition about the process, we show an example of forward substitution and reversion in the three-strain model (Figure 1). The dynamics of the simulations, where mutations occur stochastically, are compared to the analytical model by setting *t*_1_ and *t*_2_ to the times at which the strains are observed to emerge in the simulation. By analogy with the three-strain model, whichever strain that emerges first containing one mutation (either 01 or 10) is denoted strain 1. For the time interval shown here, only three strains emerge, but over longer durations, all four strains will typically be observed.

**Figure 1:**
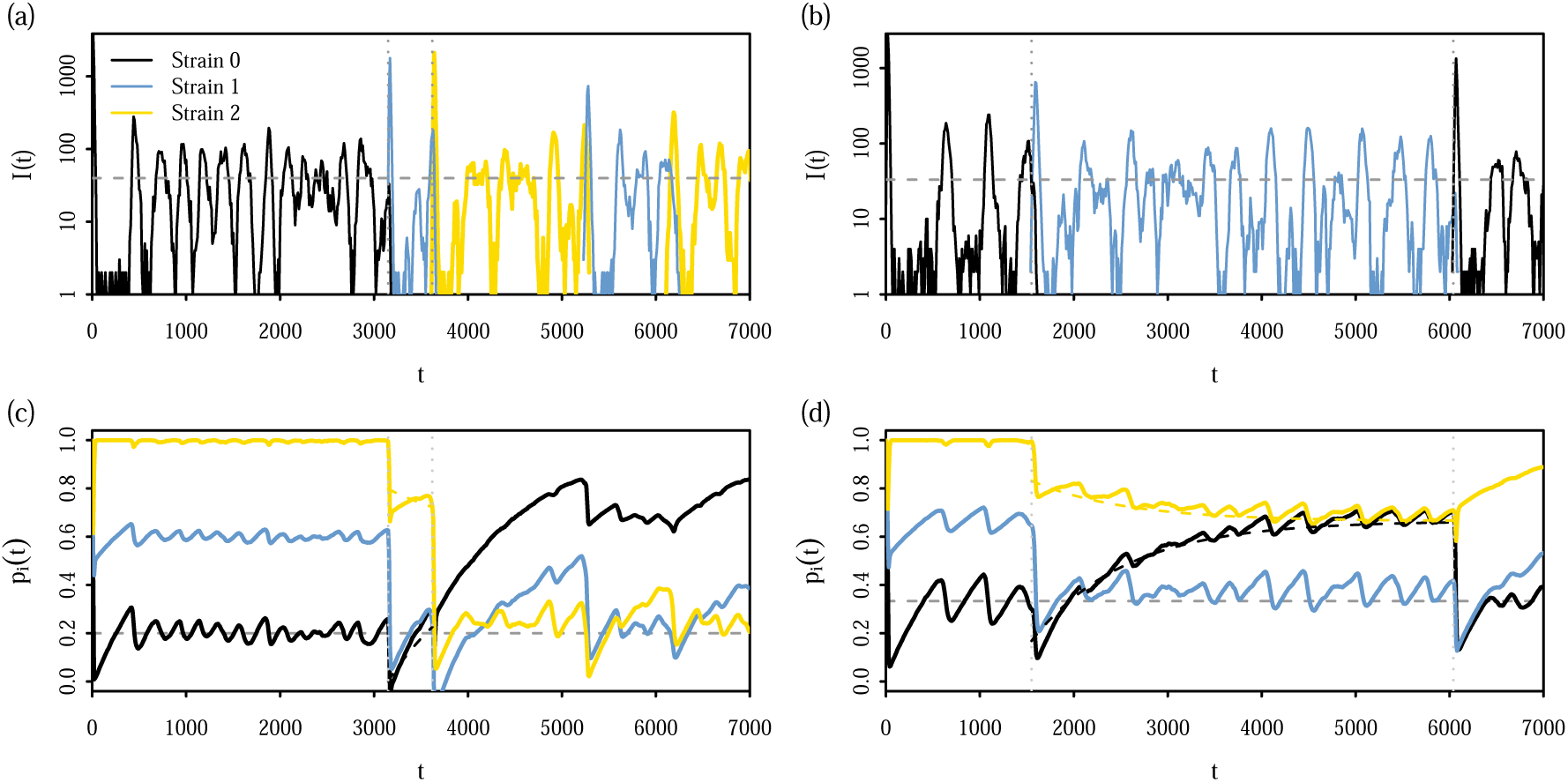
An example of forward substitution [panels (a) and (c)] and reversion [panels (b) and (d)] in the two-site bit-string epidemiological model. Solid lines show the trajectory from a single simulation of the number of hosts [panels (a) and (b)] infected by, and the proportion of hosts susceptible [panels (c) and (d)] to strains 0, 1 and 2. Simulations are initialised with a small number of hosts infected with strain 0 which tend towards the equilibrium [horizontal grey dashed lines; Equations (5–6)]. At time *t*_1_, strain 1 emerges and dominates the population until time *t*_2_ when a third strain (either strain 0 or 2) emerges. Times *t*_1_ and *t*_2_ are indicated by vertical dotted grey lines. Between *t*_2_ and *t*_1_, the expected proportion of susceptible hosts [Equations (12–15)] is shown by dashed lines. Simulations were run with parameters (a) *β* =1.0 day^−1^ and (b) *β* = 0.6 day^−1^ and in both panels, *N* = 10^4^, *δ* = 0.2, *γ* = 10^−3^ day^−1^ and *s* = 0.1.

For two separate simulations using the two-site bit-string model, we show the number of hosts infected with each strain *i* = 0,1, 2 [panels (a) and (b)], and the corresponding proportion of susceptible hosts *p_i_* [panels (c) and (d)]. The emergence of the ancestral strain 0 in the initially naive population sharply reduces the proportion of susceptible hosts to strain 0, *p*_0_; *p*_1_ is also slightly reduced due to cross-immunity between strains 0 and 1, while *p*_2_ is unaffected. When strain 1 emerges and dominates the population, both *p*_0_ and *p*_2_ are temporarily reduced but *p*_0_ slowly increases above its previous equilibrium.

In the first simulation [panels (a) and (c)], strain 1 is rapidly replaced with strain 2, so that at the time of emergence *t*_2_, susceptibility to strain 0 remains quite low [black line in panel 1(c)]. In this case, forward substitution is favoured because there is a larger pool of susceptible hosts for strain 2. In contrast, in panels (b) and (d), the interval between *t*_1_ and *t*_2_ (vertical grey lines) is longer than the first simulation, providing time for *p*_0_ to reach similar levels to *p*_2_ so that reversion can occur.

### 3.2. Time-dependence of the probability of reversion

In Figure 2, we show the probability of reversion as a function of *τ* = *t*_2_ – *t*_1_, the interval between the time of strain emergence (indicated by vertical grey lines in Figure 1). The theoretical probability of reversion [Equation (19)] is compared to the proportion of reversion events in simulations with a two-site bit-string model. We compute a proportion by binning substitution events with the same value of log_10_(*τ*), rounded to two significant figures.

**Figure 2:**
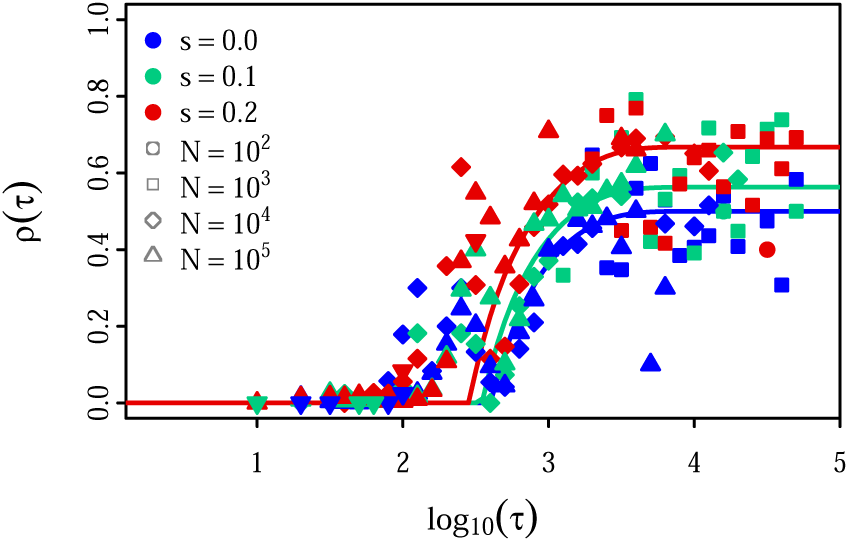
The probability of reversion, as a function of the time between strain emergence *τ*. The blue line shows the reversion probability [Equation (19)] of an unconstrained antigenic site as immunity decays, whereas the green (*s* = 0.1) and red (*s* = 0.2) lines show the combined effect of selective cost and immunity. Points show the proportion of reversion events observed from simulations of the two-site bit-string model. The proportion was computed from the binned number of substitution events that occurred immediately after a transition from strain 0 to strain 1, using the observed time between antigenic substitutions as a proxy for *τ*. Simulations were run for 10^5^ time-steps, with a time-step of one day, with 1000 replicates for each parameter combination of *s* and *N*. All other parameters were set to immune decay: *γ* = 10^−3^ day^−1^, mutation rate: *μ* = 10^−5^ site^−1^ day^−1^, recovery rate: *δ* = 0.2 day^−1^ and transmission rate: *β* = 0.6 day^−1^.

To correspond to the analytical model, only substitution events following transitions between strain 0 to strain 1 are counted. Note that in the analytical model, *τ* is the interval between the times of emergence; however, in the simulation, it is difficult to determine which of the emerging mutations will reach fixation. As a proxy for *τ*, the counts from the simulation are binned according to the time between antigenic substitutions (i.e. the time at which a different antigenic strain becomes the dominant strain in the population).

These results confirm that the reversion probability varies with *τ*. The probability of reversion is low if substitution occurs rapidly, and gradually increases with *τ* until it flattens at the asymptote *ρ*_∞_, given by Equation (21). This asymptotic value represents the probability of reversion in the absence of cross-immunity. The decay rate of host immunity *γ* affects the speed at which the asymptotic value is reached, but not the value of the asymptote.

Greater variation is seen in the simulated results for large *τ*, as these represent proportions computed from a smaller number of more rare events. However, the greatest discrepancy between theoretical and simulated results occurs near the transition *t_c_*_,0_ [Equation (18)]. At *τ* = *t_c_*_,0_, the theoretical model predicts a sharp transition away from *ρ*(*τ*) = 0; in the stochastic simulations, the transition is more gradual. The reason for this discrepancy is that the theoretical model assumes that each strain reaches equilibrium before it is replaced. However, in large viral populations, the mutational input rate can be large enough that strain 1 replaces strain 0 before *I*_0_ can reach equilibrium. In these cases, 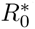 will be upwardly biased, so that *ρ*(*τ*) underestimates the probability of reversion. We confirm this in Figure B10 in Appendix B where a similar plot is shown ignoring substitution events that occur before equilibrium is reached.

Based on the form of *ρ*(*τ*), we expect the time-dependent probability to be independent of the viral mutation rate and population size. Consistent with this, we observe that simulation results for different population sizes lie on the same curve, with points from small populations (circles) corresponding to large values of *τ* and points from larger populations (triangles) corresponding to smaller values of *τ*.

### 3.3. The effect of epidemiological parameters

To examine the effects of viral transmission (*β*, *δ*, *s*) and host immunity (*γ*, *σ*), we now consider *ρ* for a fixed *τ* in the analytical model [Equation (19)]. For simplicity of notation, we omit the argument *τ* in this section. Equations (13-15) indicate that the strength of immune protection *σ* affects *ρ* only through the coefficients *A*, *B*_0_, *B*_2_, and is expected to have only a weak effect. In Figure 3, we confirm that the level of immune protection *σ* has only a weak effect on *ρ* unless the typical duration of the infection 1/*δ* [Figure 3(a)] is as long as the immune duration 1/*γ* [Figure 3(b)], or *σ* is negligibly small. Throughout the rest of the paper, we set *σ* = 1.0.

**Figure 3:**
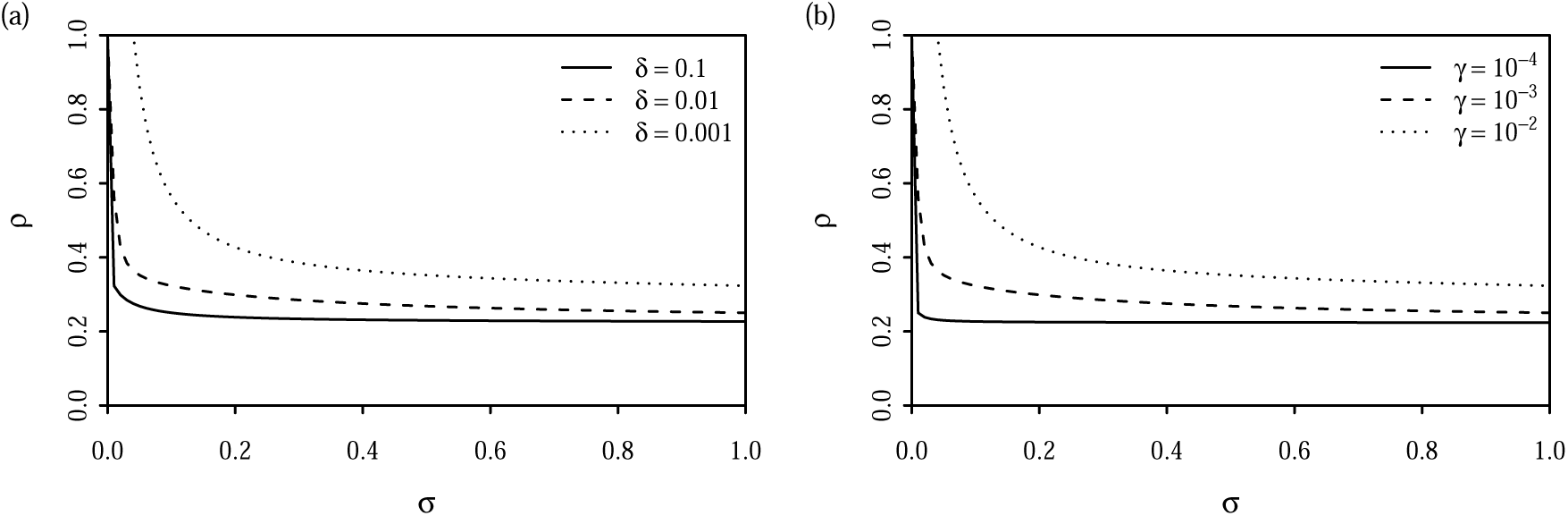
A comparison of the sensitivity of the reversion probability [Equation (19)] to the strength of immunity *σ* for different (a) rates of recovery *δ* and (b) rates of immunity decay *γ*. Unless otherwise specified, parameters were set to *γ* = 10^−3^ day^−1^, *δ* = 0.1 day^−1^, *β/δ* = 5, *γτ* = 0.5 and *μ* = 10^−5^ site^−1^ day^−1^.

Figure 4(a) shows how the reversion probability varies as a function of the basic reproductive ratio *β*/*δ*, for various values of selective cost *s*, for a fixed level of host immunity (*γτ*). For sites under no selective constraint (black line), the probability of reversion increases slightly with *β*/*δ*, but very different effects are observed for a non-zero selective cost. The effect of a selective cost is strongest for small transmission rates, as slight decreases in infection rates can have a more detrimental impact on the mutant subpopulation.

**Figure 4:**
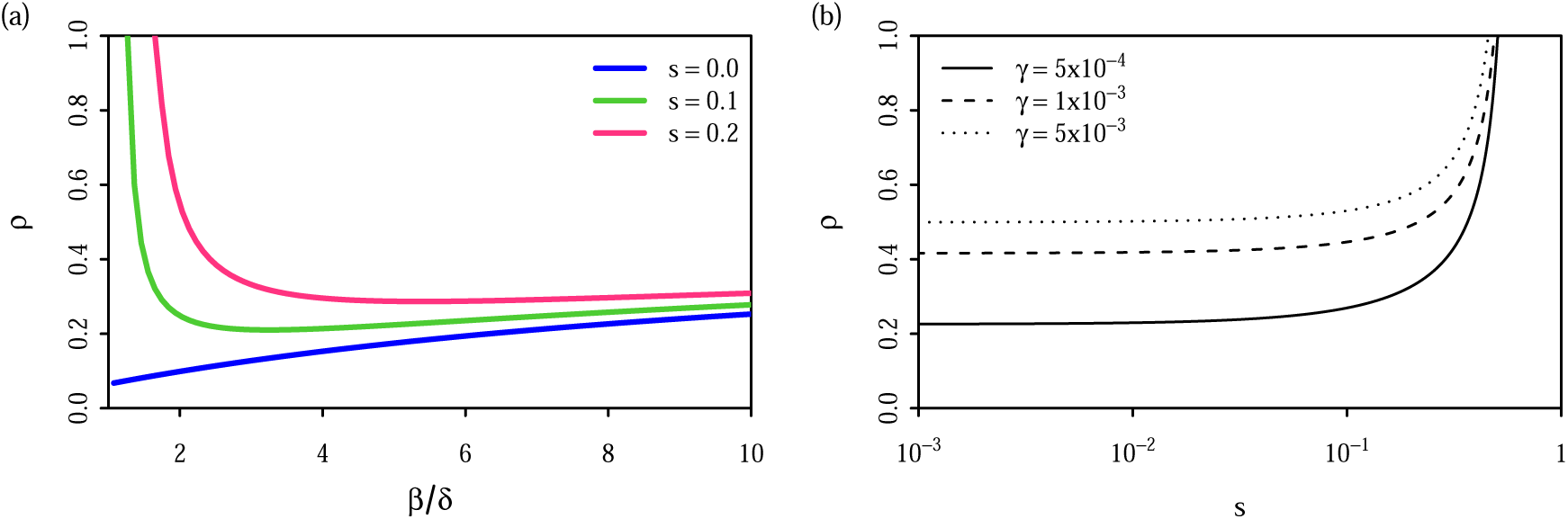
The effect of the cost of immune escape *s* and decay rate of immunity *γ* on the probability of reversion [Equation (19)]. In (a), we hold the level of immunity (*γτ* = 0.5) constant to show how varying basic reproductive ratio *β*/*δ* changes the effect of *s*. In (b), we show the effect of varying *s* for different values of *γ* with a fixed time between strain emergence *τ* = 3 × 365 days and *β*/*δ* = 5. Other parameters were set to *δ* = 0.2 day^−1^ and *μ* = 10^−5^ site^−1^ day^−1^.

The interaction between the selective cost s, and the immunity decay rate *γ*, is shown for a fixed *τ* [Figure 4(b)]. We showed in Figure 2 that for large *γτ*, *ρ*(*τ*) plateaus at *ρ*_∞_, which is independent of *γ*; however, when the rate of strain replacement is comparable to the decay rate of host immunity, there are strong dependencies. The effect of varying *γ*, in the absence of selective constraint (*s* ≈ 0), can be seen in the difference between *ρ* where the curves plateau. Further increases in selective cost leads to a rapid increase in the probability of reversion, with more rapid increases for longer lasting immunity (solid line).

### 3.4. Fluctuating frequencies at antigenic sites

In Sections 3.1-3.3, we observed that *τ* had a strong effect on whether reversions occur or not. In fact, where *τ* is known, no further information on mutation rate *μ* or population size *N* is required. However, in practice this quantity is difficult to measure. It is possible to account for variation in *τ* by integrating over the distribution of *τ*, but this can remove important information; under certain parameter ranges, the stochasticity of *τ* is sufficient to cause noticeable variation in reversion probabilities.

To observe the effect of fluctuations in *ρ*, we measure the frequency of the ancestral allele *π*_0_ at each antigenic site. The frequency of an allele is informative about its fixation probability [26], and the rate of change in frequency is proportional to the strength of selection *s* [27, 17]. Under directional selection, we expect any allele to eventually reach fixation or extinction. Thus fluctuations between *π*_0_ = 0 to *π*_0_ = 1 indicates changes in selection. We measure the frequencies of each antigenic site separately, as immunity against each site may vary depending on the history of previous circulating strains.

In Figure 5, we show frequency trajectories *π*_0_, under conditions of both antigenic selection and selective constraint, so that antigenic changes away from the ancestral sequence imposes a cost. To account for inaccuracies due to sampling, *π*_0_ was computed from sequences sampled at discrete intervals, and the earliest sequence sampled after the burn-in period was used as the ancestral sequence. In all panels, we observe fluctuations in frequency levels as reversion probabilities vary due to the stochasticity of the time between antigenic substitutions, although there is no change in *μ*, *N*, or *s* during a simulation. The pattern of fluctuations in *π*_0_ differs depending on the host population size *N* (varying along columns) or the decay rate of host immunity *γ* (varying along rows). Faster changes in *π*_0_ are observed for larger *N* and fixation of the ancestral allele becomes less likely. Tracking frequency over time also provides information on *γ* that would not be available in the time-averaged approach. Comparison between columns in Figure 5 indicates that increasing *γ* tends to reduce both the frequency and amplitude of *π*_0_. This effect is particularly evident for larger population sizes [panels (c)–(f)], where the rate of substitution is not limited by the rate of mutational input.

**Figure 5:**
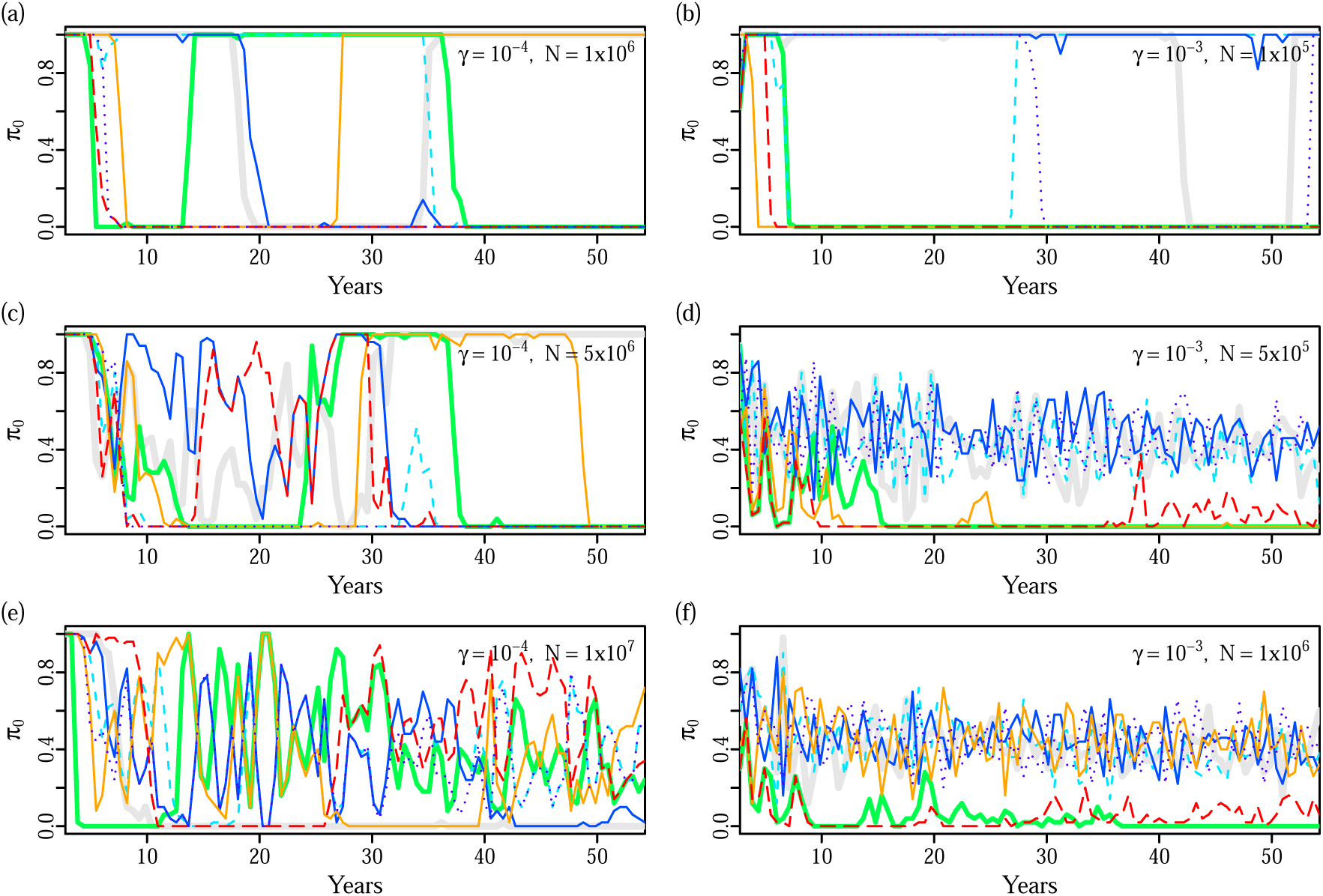
The effect of population size and duration of immunity on the frequency of the ancestral allele *π*_0_ at antigenic sites under selective constraint (*s* = 0.2). Each line represents the changing frequencies, at a single antigenic site, of the ancestral allele estimated to be the earliest sampled amino acid residue after the burn-in period (1000 days). For high rates of mutational input [panels (d) and (f)], the earliest sequence may not be the true ancestral sequence (set to be the most transmissible), which in some cases results in low observed values of *π*_0_. Each panel represents the dynamics of a single simulation, with *π*_0_ computed from samples of 20 sequences taken every 200 days. All simulations were run with a time-step of one day and parameters *β* =1.0 day^−1^, *δ* = 0.2 day^−1^, *μ* = 10^−5^ site^−1^ day^−1^, *L_a_* = 7, to match parameters used for human influenza A (H3N2) [43].

The effect of removing the selective cost (*s* = 0) is shown in Figure 6. Although fluctuations can still occur, the ancestral allele at the antigenic site rarely returns to fixation (*π*_0_ = 1) and, if so, does not remain fixed for long. This effect occurs even for small population sizes [panel (a)] which favour reversion. Continual antigenic selection drives further substitutions to other derived amino acid residues, that have not induced prior immunity. That is, multiple instances of increasing *π*_0_ as an indication of high selective costs *s* is robust to misspecification of the ancestral allele. However, consistently low values of *π*_0_ may simply be due to using an misspecified ancestral allele (an alternative interpretation is that *π*_0_ correctly identifies that an unfavoured amino acid is unconstrained).

**Figure 6:**
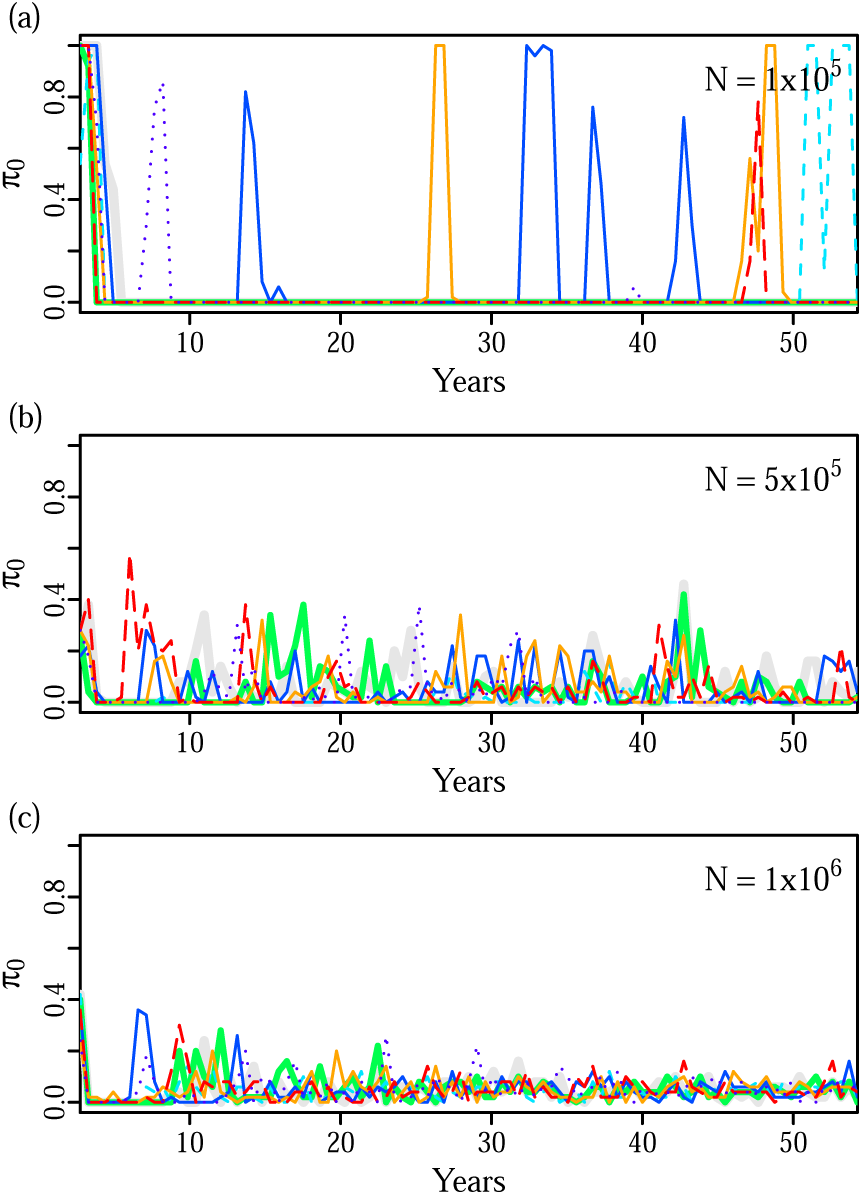
The frequency of the the ancestral allele *π*_0_ at antigenic sites under no selective constraint (*s* = 0). Lines in each panel show the changing frequency of the ancestral (earliest sampled) allele at each antigenic site in a single simulation. *π*_0_ was computed from samples of 20 sequences taken every 200 days, discarding all sequence data from the burn-in period of 1000 days. All simulations were run with *γ* = 1 × 10^−3^ day^−1^, *β* = 1.0 day^−1^, *δ* = 0.2 day^−1^, *μ* = 10^−5^ site^−1^ day^−1^, and *L_a_* = 7.

### 3.5. Application to influenza and RSV

In Figure 7, we show *π*_0_ changing over time for the human influenza A virus subtype H3N2 and the respiratory syncytial virus (RSV) subtype A at antigenic and non-antigenic sites. The H3N2 data set consists of all HA sequences for human H3N2 from the influenza virus database [28] where the year of sampling is known. The accession numbers surface G protein sequences of RSV-A sequences that we used were listed in Botosso et al. [5]. In total, we analysed 5831 H3N2 sequence spanning 45 years and 538 RSV sequences spanning 19 years.

**Figure 7:**
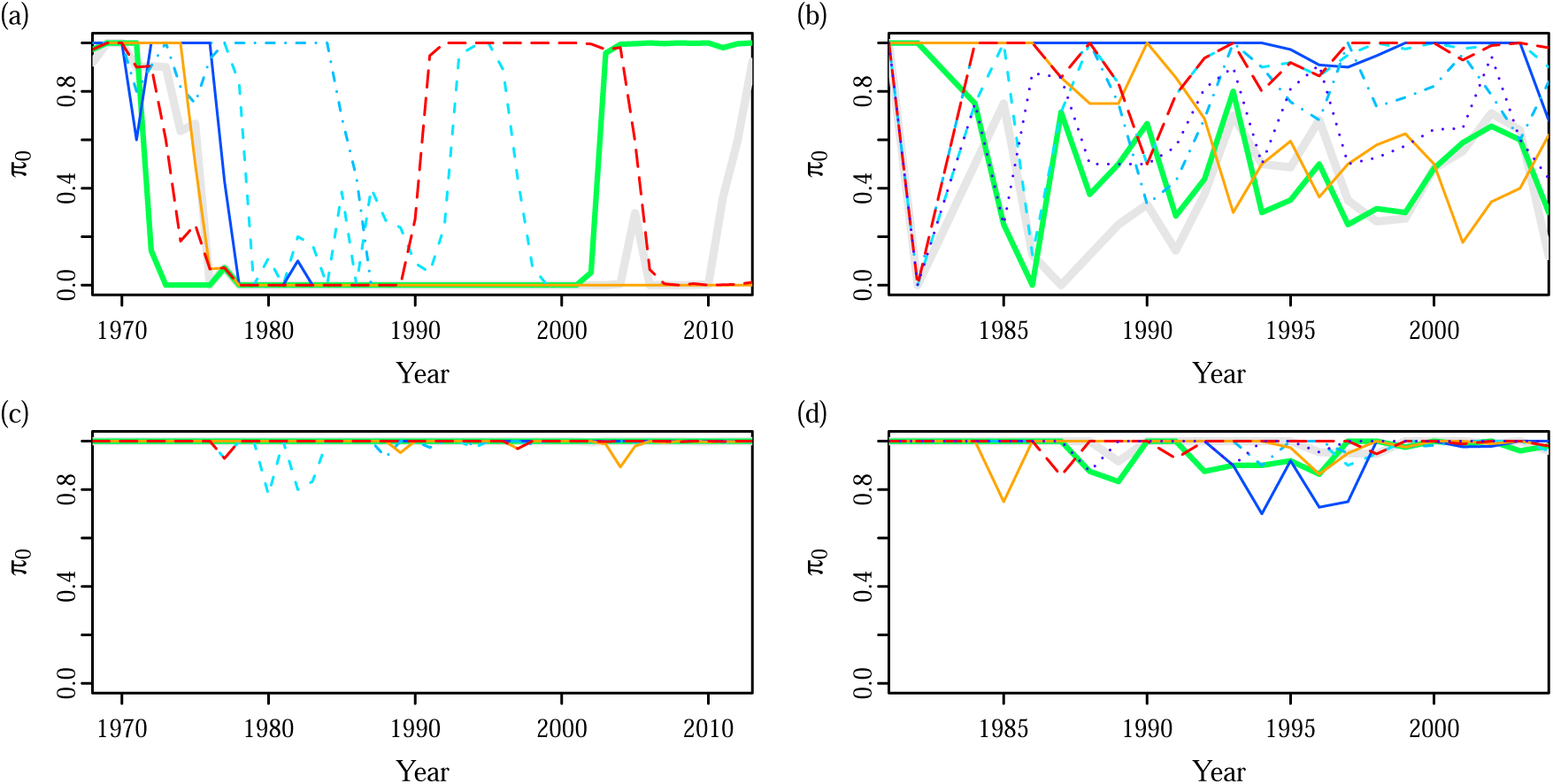
Trajectory of the frequency of the ancestral allele *π*_0_ computed at antigenic sites of (a) human H3N2 and (b) human RSV-A show fluctuations which are distinct from randomly chosen non-antigenic sites [panels (c) and (d)]. Frequencies were computed at (a, c) seven sites in the HA segment of H3N2 with A/Aichi/2/1968 as the ancestral strain, and (b, d) eight sites in the C-terminal hyper-variable region of the surface G protein of RSV-A using strain AF065406 (sampled in 1981) as the ancestral strain. Sequences were pooled according to the year of isolation, with years in which fewer than five sequences were sampled were excluded.

We computed *π*_0_ for antigenic sites which have been identified by experimental methods, as sequence-based methods are also designed to identify sites with variation in amino acid composition. For H3N2, we used the seven sites (145, 155, 156, 158, 159, 189, 193) listed in a recent study [29] which used antigenic cartography which integrates information over multiple pairs of antigen and antisera in order to evaluate overall antigenic change [30]. For RSV-A, experimental studies with monoclonal antibodies have identified a large number of sites which react to different monoclonal antibodies [31, 32]. More recent studies have used phylogenetic analysis of natural isolates to identify potential antigenic sites [33, 34]. Note that there is an ascertainment bias in using sites identified on the basis of frequent amino acid changes. Here, we have restricted the analysis to eight sites (225, 226, 233, 237, 244, 274, 280, 290) which were identified as reducing antigenic recognition in multiple studies [33, 34, 31, 32], with at least one being experimental [31, 32]. Including a larger number of sites does not affect the results, but will obscure features of distinct trajectories.

For both viruses, we obtain oscillating patterns of *π*_0_ that are consistent with our expectations for antigenic sites evolving under both immune selection and functional constraint. Non-antigenic sites [Figure 7(c) and (d)] generally do not exhibit these fluctuations, but some non-antigenic sites in RSV-A may experience frequency fluctuations due to linkage to antigenic sites [Figure 7(d)]. Patterns of frequency change in H3N2 and RSV-A differ considerably from each other. H3N2 frequencies have sharper and slower oscillations, which are suggestive of both a smaller population size and longer lasting immunity. At least four antigenic sites in H3N2 revert and fix at the ancestral state which indicates very strong selective constraint. RSV-A shows more rapid oscillations, suggesting faster decaying immunity and moderate selective constraint. The relatively short time that the ancestral allele is at high frequencies suggests that selective constraint has a smaller influence than for H3N2.

## 4. Discussion

We have shown that for acute, recurrent infections, the probability of reversion at antigenic sites depends on the interaction between the cost of immune escape and the duration of host immunity. Similar to models for HIV [17], we find that a higher cost of immune escape increases the probability of antigenic reversion. The impact of the cost of immune escape on the reversion probability is greater when the basic reproductive ratio is low, as small reductions in transmissibility have a more detrimental effect. This is in agreement with a previous study on the effect of selective constraint on antigenic drift [35]. In addition to these two parameters, we find that longer lasting immunity can also reduce the probability of reversion, but the precise extent of this reduction depends on the time between antigenic substitutions.

The time between antigenic substitutions, which is inversely proportional to the viral population size and mutation rate, is closely related to the rate of mutational input *θ*, a parameter commonly used in population genetics to describe the time-scale of selection and drift. In the epidemiological model, it affects the balance between selective constraint and antigenic selection by determining the extent to which prior immunity has decayed. When the interval between antigenic substitutions is small, immunity against the ancestral strain remains high at the time of substitution so that antigenic selection reduces the reversion probability. For larger intervals between antigenic substitutions, prior immunity will have decayed to a greater extent and the basic reproductive ratio and cost of immune escape become stronger determinants of the reversion probability. In the context of phylodynamic models, *θ* is also the parameter which is used to link the coalescent to epidemiological models [36].

Previous studies have described varying levels of reversion in a range of viruses and speculated on the influence of host immunity [5, 2, 8], but it has been unclear how the level of reversion should be quantified and how these results should be interpreted. In contrast to previous studies [5, 37] based on phylogenetic methods, we propose using temporal patterns of frequency change to quantify reversion. Where sequence data from multiple time-points is available, a frequency-based approach can more easily show the time-dependent effect of antigenic selection. Simulation results predict that varying parameters controlling population size, transmission rate, immunity decay and selective constraint have qualitative effects on the frequency of the ancestral allele *π*_0_ which are consistent with the analytical model, providing a means for interpretation. As our approach uses site-frequency data rather than a phylogeny, it is amenable to the application of large time-structured data sets, but is also more sensitive to effects such as biased sampling and spatial structure.

In this paper, we compared patterns of *π*_0_ for two viruses that induce acute respiratory infection which recurrently infect human populations and induce long-term immunity: influenza A (H3N2) and RSV-A. For both viruses, we observed fluctuations in frequency at antigenic sites suggesting the presence of both immune memory and selective constraint. Without the continuously changing balance between these two effects, we would expect an allele for a particular site to reach fixation and remain in that state [27]. While RSV has been reported to experience high levels of reversions [5], previous phylogentic studies have not identified reversion in H3N2. However, a recent study [8] showed that changes at antigenic sites in H3N2 occur as cycles in a genotype network; that is, mutations to multiple states occur before reversion to the ancestral allele, so that the reversion is not identifiable along a the phylogeny.

Our model suggests that the higher rates of reversion in RSV-A compared to H3N2 is due mainly to more rapidly decaying immunity rather than stronger selective constraint. Fluctuations in frequency are more rapid and complete fixation of the ancestral amino acid does not occur for most antigenic sites of RSV-A. In contrast, for H3N2, we observe multiple occasions where a fixation of the ancestral amino acid occurs, and long periods where *π*_0_ = 1 is maintained, suggesting strong selective constraints. This is consistent with the location of the sites within the receptor binding region of the HA gene, so that any antigenic change is also likely to affect viral transmissibility [29]. Comparison between the frequency of the oscillations also suggests that H3N2 induces more long-lasting immunity than RSV-A. RSV-A exhibits more rapid fluctuation while several of the antigenic sites in H3N2 were fixed for long periods (> 10 years) at a derived amino acid, supporting the hypothesis that immune pressure against reversion is maintained for long periods. Frequency patterns of H3N2 frequency patterns are consistent with multi-site codon simulations (Figure 5) with host immunity decay rate *γ* on the order of 10^−4^, whereas a value of *γ* ≈ 10^−3^ is more compatible with frequency patterns for RSV-A. These values are in agreement with reinfection experiments which estimate immunity for H3N2 lasting 8 years (*γ* = 3 × 10^−4^ day^−1^) [38] compared to 1.8 years (*γ* = 1.5 × 10^−3^ day^−1^) for RSV-A [39].

Our study shows that the frequency of the ancestral allele, *π*_0_, which can be easily calculated for time-stamped viral sequences, is informative about the immune dynamics and cost of escape. In particular, sharp fluctuations in frequency is indicative of immune selection occurring at a comparable time-scale to substitutions at antigenic sites. However, a small number of linked sites may also display similar patterns as they co-segregate with antigenic sites. That is, frequency patterns should not be used as a method to identify antigenic sites; but where the antigenic sites are known, frequency patterns provide information about the epidemiology of the virus as a whole.

The approach outlined here provides a qualitative description rather than estimates of the epidemiological parameters. Analytical expressions, relating the probability of reversion to the parameters underlying the viral dynamics for the three-strain model, rely on the assumption that each strain reaches equilibrium before it is replaced. This assumption tends to be violated when population sizes and mutation rates become large, so that we generally underestimate the probability of reversion. To address the restrictions of the equilibrium assumptions and the assumption of only three strains, we used computer simulations describing sequence dynamics in a multi-site model. Formal inference using a complex computational model is a challenge for future research. Despite the simplicity, our approach is useful in providing a scheme to consider both epidemiological and molecular effects simultaneously. As such, it is complementary to both coalescent approaches [40, 41, 42] which assume epidemiological dynamics are largely unaffected by molecular changes, and to codon-based methods [14, 15] which assume that substitution occurs instantaneously as a time-homogeneous process along branches of the phylogeny.

Our model highlights the importance of understanding the interaction between epidemiological and molecular effects. The results imply that different evolutionary trajectories are expected in viral populations with the same distribution of fitness effects but differing population size and contact rates. In particular, we expect that viral populations in larger cities with denser populations undergo less reversion and are more likely to generate antigenically novel variants.

## Acknowledgements

We thank Peter White for discussions about reversion in viruses. This work was supported by a Discovery Grant DP110100465 from the Australian Research Council and by the Australian Postgraduate Award to CHSC.

## Appendix A. Comparison of the compartmental SIRS model with an agent-based model for a single strain

The compartmental SIRS model described in Section 2.1 tracks only the number of hosts with immunity in the population *R*, which is increased with each infection by an increment of *σ*. The model, however, does not account for the fact that partially immune hosts which are re-infected cannot gain more than complete immunity. To examine the effect of this approximation, we implement, for the single strain case, an agent-based model which tracks the level of immunity *r_i_* ∈ [0,1] in each host *i* in the population. The agent-based variables can be related to the population model [Equation (3)] by summing across all uninfected hosts 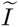,

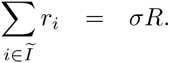

We implement two agent-based simulations which differ in how they model the decay of host immunity. In model A1, immunity decays deterministically, so there is no variability in the rate of decay between hosts. In contrast, in model A2, we maximise the variability in the rate of decay by having complete loss of immunity in a proportion of hosts. In both models, population-wide levels of immunity are reduced, but there are considerable differences in the variation in levels of immunity between hosts, which is known to affect the epidemiological dynamics [44].

1. Transmission: The number of new infections in each time-step is Poisson with mean *βSI/N*. Newly infected hosts are randomly drawn with the set of uninfected hosts by multinomial sampling, according to 1 – *r_i_*. Note that their immunity status is not altered on infection, so they retain immunity obtained from prior infections, but their contribution is not included into the population variable *R*.
2. Recovery: The number of infected hosts which recover in each time-step is Poisson with mean *δI* truncated with an upper bound of *I* – 1. Each recovered host *i* is drawn from the infected population with uniform probability and their immunity is increased by

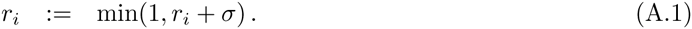
3. Decay of host immunity is simulated differently between models A1 and A2. In model A1, immunity is reduced deterministically in each uninfected hosts *i*

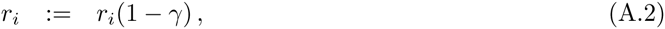

whereas in model A2,

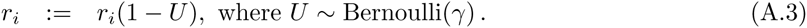

We compare the agent-based models to the compartmental model implemented in Section 2.2 with no mutation (single-strain). Importantly, the equilibrium value for *S* is largely unchanged between models [Figure A.8(b)], suggesting that derivations of *ρ*(*τ*) remain valid. However, differences can be seen in equilibrium value of *I*. The compartmental model equilibrates at a lower mean value of *I* compared to both agent-based models A1 and A2, with a larger difference when compared to A1 (Figure A.9). However, this discrepancy is comparable to the variance in the of *I*(*t*) over time [Figure A.8(a)]. The difference in *I* between simulations is largely unaffected by changes in population size *N* or transmission rate *β*, but mainly influenced by *σ* (Figure A.9).

**Figure A.8:**
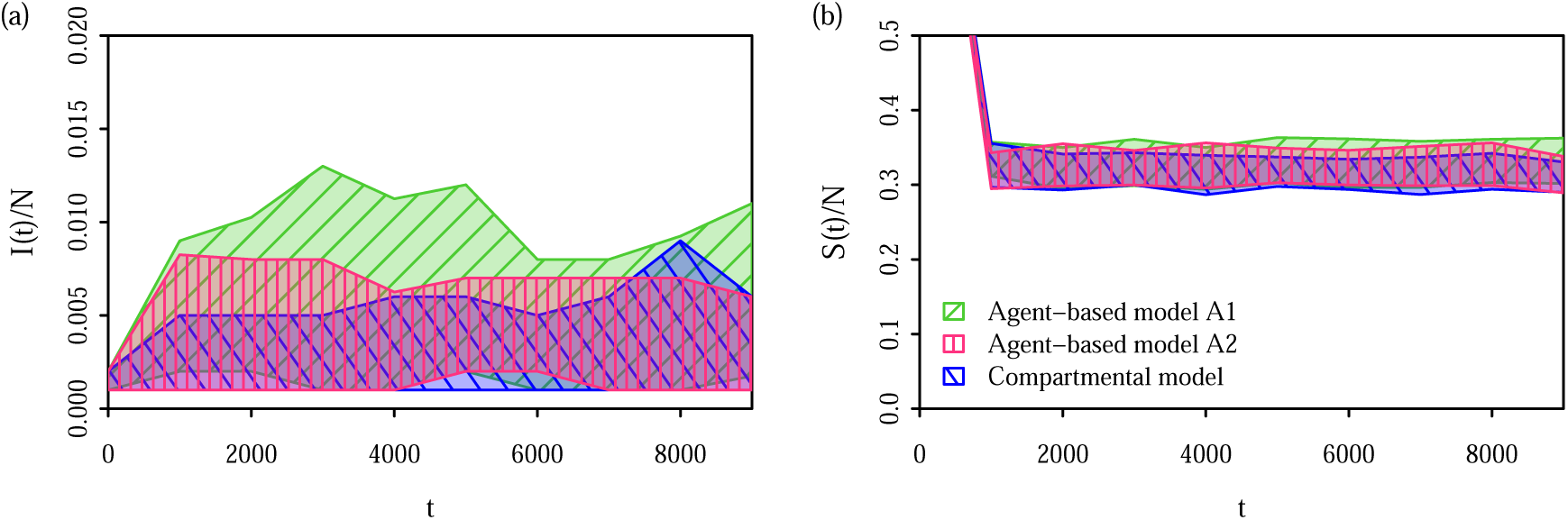
Comparison of the simulation dynamics between agent-based and compartmental simulations. Shaded areas show the interquartile range for 100 replicates with *β* = 0.6 day^−1^, *δ* = 0.2 day^−1^, *γ* = 10^−3^ day^−1^, *σ* = 0.8, *N* = 10^3^.

**Figure A.9:**
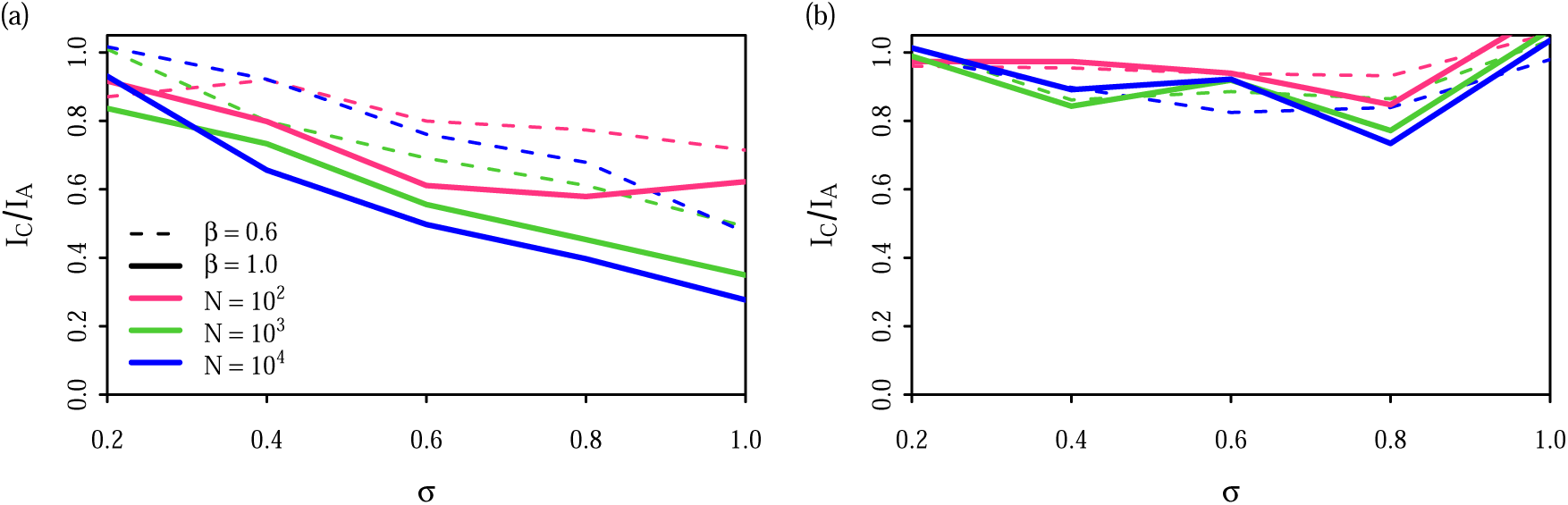
Underestimation of the size of the infected population in the compartmental model compared to the agent-based models (a) A1 with gradual decay of host-immunity and (b) A2 with sudden decay of host-immunity. The extent of underestimation was computed by taking the mean of *I*(*t*) over 100 simulations over 10^4^ time-steps for both the compartmental model and the agent-based model, and taking the ratio of the means. Simulations were run with *δ* = 0.2 day^−1^ and *γ* = 10^−3^ day^−1^.

**Figure B.10:**
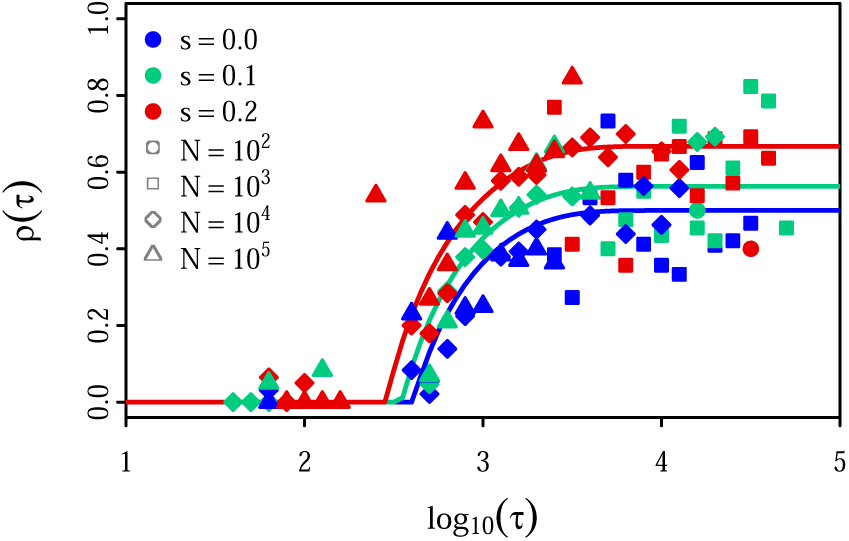
The probability of reversion, as function of the time between strain emergence *τ.* This is the same as Figure 2, except that substitution events which occur before the population reaches equilibrium are ignored.

## Appendix B. Effect of equilibrium assumption in SIRS model

The derivation of the probability of reversion [Equation (19)] depends heavily on the assumption that the system reaches equilibrium before antigenic substitutions occur. This effect can be seen by comparing Figure 2 to Figure B.10 where substitution events that occur before equilibrium is reached are ignored. The points with the greatest discrepancy to the theoretical values of *ρ*(*τ*) for intermediate values of *τ* are are not seen in Figure B.10. There is more variation from theoretical expectations for large *τ*, as more substitution events have been removed the calculation.

